# A framework for estimating the effects of sequential reproductive barriers: implementation using Bayesian models with field data from cryptic species

**DOI:** 10.1101/363168

**Authors:** Jean Peccoud, David R. J. Pleydell, Nicolas Sauvion

## Abstract

Determining how reproductive barriers modulate gene flow between populations represents a major step towards understanding the factors shaping the course of speciation. Although many indices quantifying reproductive isolation (RI) have been proposed, they do not permit the quantification of cross direction-specific RI under varying species frequencies and over arbitrary sequences of barriers. Furthermore, techniques quantifying associated uncertainties are lacking, and statistical methods unrelated to biological process are still preferred for obtaining confidence intervals and p-values. To address these shortcomings, we provide new RI indices that model changes in gene flow for both directions of hybridization, and we implement them in a Bayesian model. We use this model to quantify RI between two species of the psyllid *Cacopsylla pruni* based on field genotypic data for mating individuals, inseminated spermatophores and progeny. The results showed that pre-insemination isolation was strong, mildly asymmetric and undistinguishably different between study sites despite large differences in species frequencies; that post-insemination isolation strongly affected the more common hybrid type; and that cumulative isolation was close to complete. In the light of these results, we discuss how these developments can strengthen comparative RI studies.

**Author contributions:** JP and NS initiated the study and obtained biological data. JP and DRJP developed the porosity-based approach. DRJP conceived the Bayesian implementation and code. JP, DRJP and NS wrote the manuscript.

**Data availability:** Mitochondrial sequence data will be available at Genbank, source code is available at xxx.

## Introduction

Speciation involves the build-up of reproductive isolation (RI) at several key parts of the populations’ life cycles, which are referred to as reproductive barriers. Understanding how these barriers act in conjunction to reduce gene flow and permit the divergence of populations into species has been an important goal of speciation research (Coyne and Orr 2004; Sobel et al. 2010; Butlin et al. 2012). As a result, the last fifteen years have seen a burgeoning of methods to estimate the strength of reproductive barriers from field data on natural populations (Ramsey et al. 2003; Malausa et al. 2005; Martin and Willis 2007; Matsubayashi and Katakura 2009; Sanchez-Guillen et al. 2012; Sobel and Streisfeld 2015; Pombi et al. 2017). Field estimates indeed provide the most pertinent results to help identify local factors affecting the course of speciation (Nosil et al. 2009; Via 2009; Sobel et al. 2010; Butlin et al. 2012).

This objective requires that RI estimates represent evolutionary relevant quantities, mainly potential gene flow, whilst correcting for differences in species frequencies that do not reflect phenotypic variations – effects we collectively refer to as “contingency”. In the progression toward this goal, many indices to quantify RI have been developed [reviewed in Sobel and Chen (2014)]. Key developments include formulas to quantify cumulative RI over sequential reproductive barriers (Ramsey et al. 2003); corrections for unequal species frequencies and allochrony (Martin and Willis 2007); and the integration of these developments into RI indices that maintain a linear relation to the probability of gene flow – a desirable property when comparing populations and species (Sobel and Chen 2014).

Despite this growing diversity and sophistication of RI indices, and of the studies using them, two deficiencies of current methods remain apparent. First, although RI is commonly asymmetrical (e.g., Lowry et al. 2008; Matsubayashi and Katakura 2009; Sanchez-Guillen et al. 2012; Brys et al. 2014; Kaufmann et al. 2017; Martin et al. 2017), we lack indices that can estimate directional (cross-type specific) RI over an arbitrary combination of reproductive barriers, while controlling for contingency. Second, speciation research would benefit from more studies reporting and discussing uncertainty in RI (e.g., Merrill et al. 2011; Lackey and Boughman 2017). The RI literature is dominated by the discussion of point estimates for which there exists a lack of associated uncertainty measures. Thus, it remains difficult to demonstrate whether apparent differences in RI (observed between different barriers, sub-populations or species) reflect real phenotypic differences or merely sampling error. Accordingly, practitioners seeking a richer statistical analysis - involving confidence intervals or significance tests for example - have been constrained to adopt less biologically-motivated indices such as those provided by generalized linear models (Takami et al. 2007; Polacik and Reichard 2011; Falk et al. 2012; Peccoud et al. 2014; Kostyun and Moyle 2017).

Why a more complete statistical framework for RI estimation has not emerged may partly stem from the fact that the calculation of RI is frequently complexified by the need to correct for contingency and the effects of reproductive barriers not under scrutiny, or to combine the effects of several barriers (Sobel and Chen 2014). Accounting for asymmetry in RI would further complexify existing formulas and pose a substantial challenge regarding the construction of confidence intervals and significance tests for these indices.

We suggest that these issues can be resolved by focusing attention on estimating the probabilities of gene flow – rather than RI *perse* – induced by both within- and between-species crosses. Focusing on the probabilities of gene flow facilitates statistical estimation, from field data, of contingency-independent RI indices (in both cross directions) at any reproductive barrier or over any arbitrary sequence of barriers. Moreover, this approach naturally lends itself to Bayesian uncertainty analysis. In other branches of ecology and evolution, Bayesian techniques have long been popular for numerous reasons, including: they provide a natural paradigm to account for multiple sources of uncertainty; they facilitate the incorporation of prior knowledge; they are applicable to a wide variety of models; and inference based on posterior distributions of model parameters is easy and intuitive (Gelman et al. 1995; Clark 2005; Cressie et al. 2009; Beaumont 2010; Hoban et al. 2012; Gompert et al. 2017). We illustrate these benefits with a Bayesian model designed to quantify the weight of evidence for spatial heterogeneity in RI using genetic data from natural populations of the psyllid *Cacopsylla pruni*.

## Methods

### Modeling sequential reproductive barriers

Consider two species *A* and *B* interacting at a reproductive barrier. Let *G_XY_*denote the probability that an individual sampled from the next generation comes from an *X*× *Y* cross (with *X*, *Y* ∈ {*A*,*B*} and the maternal species always noted first) in the absence of further isolation after the barrier. Thus, *G_XY_* is the potential gene flow (which we may simply call “gene flow” afterwards) induced by *X*×*Y* crosses. Let ***G***={*G_AA_*, *G_BB_*, *G_AB_*, *G_BA_*} be the set of all such proportions, which sum to one.

Estimating RI as the decrease of interspecific gene flow (Sobel and Chen 2014) requires a measure of gene flow that is independent of contingency. We call these contingency-independent gene flow rates “barrier porosities” to convey that they solely depend on phenotypic differences expressed at the barrier. We denote barrier porosities as ***β***={*β_AA_*, *β_BB_*, *β_AB_*, *β_BA_*}, these sum to one and each *β_XY_* element equals 1/4 in the absence of RI.

The ratio of porosity, over its null expectation when RI=0, indicates the strength of RI at a barrier (Sobel and Chen 2014). Thus, a bidirectional RI index that considers both hybrid cross-types is

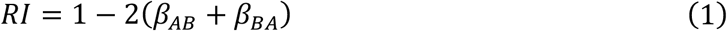

and the RI affecting just one hybrid cross-type (*X* ≠ *Y*) is:

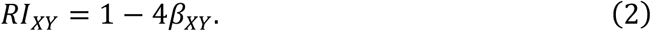

These indices vary linearly with gene flow, take value zero when porosities are 1/4, and take value one when porosity to hybridization is zero.

Directional RI indices allow between cross-type differences (asymmetry) in RI to be quantified as

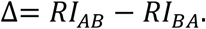

Given the simplicity of these developments, the main modeling task is to establish the relationships between gene flow ***G*** and barrier porosities ***β***. To this aim, we introduce “null gene flow” E_0_[*G_XY_*] to denote gene flow in the absence of RI at the studied barrier. E_0_[*G_XY_*] can be visualized as the flow of genes going through the previous barrier and arriving at the focal barrier. Table 1 provides examples of E_0_[*G_XY_*] and *G_XY_* for different sources of RI.

**Table 1.** Formulations for null (in the absence of RI at a studied barrier) gene flow due to females of species *X* and males of species *Y* (E_0_[*G_XY_*]), and for potential gene flow following modification by the barrier (*G_XY_*).

| source of RI | $E_0[G_{XY}]$ | $G_{XY}$ |
| --- | --- | --- |
| ecological niche difference | $f_X \times m_Y^\dagger$ | p.‡ of encounters involving females of species $X$ and males of species $Y$ (a) |
| allochrony | (a)<sup>§</sup> | p. of encounters between sexually active mates involving females of species $X$ and males of species $Y$ (b) |
| conspecific mate preference | (b) | p. of mating pairs involving females of species $X$ and males of species $Y$ (c) |
| mechanical incompatibility | (c) | $f_X \times$ (average proportion of species $Y$ sperm per female of species $X$) (d) |
| gametic incompatibility | (d) | p. of zygotes from females of species $X$ and males of species $Y$ (e) |
| early hybrid mortality | (e) | p. of progeny from females of species $X$ and males of species $Y$ (f) |
| late hybrid mortality | (f) | p. of older progeny from females of species $X$ and males of species $Y$ |

Barriers are shown in their order of appearance in the life cycle. $^\dagger f_X$ : proportion of species $X$ in females; $m_Y$ : proportion of species $Y$ in males. $^\ddagger p$ .: proportion. $^\S$ Letters in parentheses correspond to formulations proposed for $G_{XY}$ in the rightmost column. Formulations may be amended depending of the biological model, for instance individuals may be interpreted as gametes (e.g., pollen for males), and proportion of encounters can be estimated from range or habitat overlap.

At the postzygotic level, the relative frequency of *XY* genotypes in the progeny, *G_XY_*, is proportional to their frequency before the barrier, E_0_[*G_XY_*], multiplied by their probability to survive (or pass) through the barrier (defined as *S_XY_*):

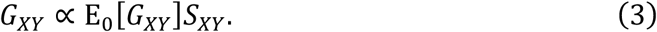

Under equal species frequencies and random mating, *G_XY_* = *β_XY_* by definition and E_0_[*G_XY_*] = 1/4. Assuming that survival *S_XY_* is constant, and is therefore independent of species frequencies, the above equation translates to:

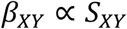

where 1/4 was dropped as a constant in the proportionality relationship.

Substituting *S_XY_* in equation 3 yields:

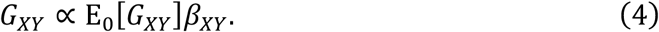

Postzygotic barrier porosities are therefore proportional to the ratio of potential over null gene flow – a metric that enables RI quantification when null expectations (elements of E_0_[***G***]) are unequal [see appendix D of Sobel and Chen (2014)].

For total gene flow to equal one, equation 4 requires normalization:

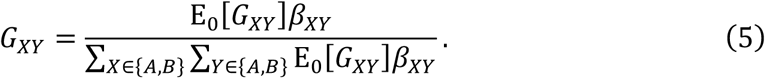

Equation 5 satisfies that when all null gene flows equal 1/4 (equal species frequencies), the porosity *β_XY_* equals gene flow *G_XY_*. Conversely, in the absence of RI (*β_XY_* is 1/4 for all combinations), gene flows equal null gene flows.

Given estimates of gene flow before and after a barrier, the porosities can be recovered by rearranging and normalizing equation 4 so that element of ***β*** sum to 1, hence:

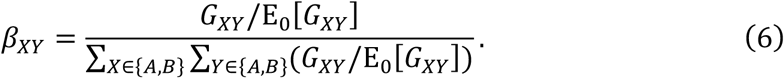

Contrarily to postzygotic barriers that increase progeny mortality, prezygotic barriers do not usually incur a fitness cost to parents. Hence, prezygotic isolation must be modeled in such a way that it does not directly affect fitness. To do so, we express the proportion of *XY* zygotes (that we expect if no isolation exists after the studied barrier) among those having a mother from species *X*:

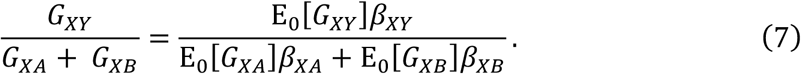

In order to derive *G_XY_*, the relative contribution of species *X* females to the next generation (*G_XA_* + *G_XB_*) must be specified. If, at the focal barrier, female reproductive success does not vary between the species, then *G_XA_* + *G_XB_* is the frequencies of species *X* in females, which we call *f_x_*, and equation 7 becomes:

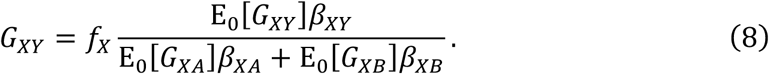

Barrier porosities can be obtained from gene flows by rearranging equation 7 into (proof not shown):

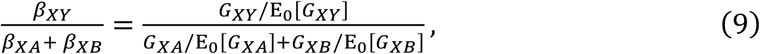

and specifying *β_XA_* + *β_XB_* appropriately. If female reproductive success can be assumed equal between species, then *β_XA_* + *β_XB_* = 1/2, so:

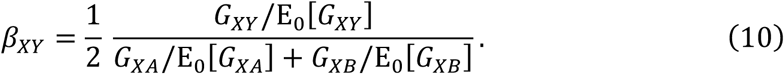

Equation 7, and its by-products, assume that the fraction of *XY* zygotes contributed by females of species *X* is proportional to its null-expected value in the absence of RI, E_0_[*G_XY_*]. This implies that the probability of hybridization per interspecific encounter, represented by the ratio *G_XY_*/E_0_[*G_XY_*], does not depend on species frequencies, and reflects the barrier porosity *β_XY_*. If the assumption of equal reproductive success between females is not warranted, alternative formulations for equations 8 and 10 may be desirable. Such developments should be tailored to the specifics of the biological system and are beyond the scope of the current work. Once obtained, barrier porosities can be used to model a sequence of *b* barriers with porosities ***β***^1^ … ***β***^*b*^. The product of these porosities is proportional to the probability that genes from two parents flow through all these barriers to eventually produce an offspring. The combined porosity of these barriers to *X*×*Y* gene flow is thus given by:

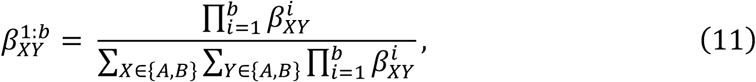

whose denominator ensures that the combined porosities of all four *XY* combinations sum to one.

Equations 5 and 8 permit barrier porosities *β_XY_*, hence RI, to be estimated via statistical techniques that confront modelled gene flows *G_XY_* with data collected at different points of the reproductive cycle. We will demonstrate this approach with a Bayesian implementation. Alternatively, a simpler approach would use equations 6 or 10 to obtain point estimates of barrier porosities by specifying *G_XY_* according to observations (examples given in Table 1).

### Study model

Our model system, *Cacopsylla pruni* Scopoli (Sternorrhyncha: Psyllidae), includes two unnamed cryptic species which are strongly genetically divergent but have yet to show ecological or morphological differences (Sauvion et al. 2007; Peccoud et al. 2013). These species co-occur at several sites in Southern France (Sauvion et al. 2007) on shrubs of genus *Prunus*, on which the insects feed, reproduce and die in spring (Figure 1). Progeny reach adulthood after approximately 2 months, migrate shortly after to conifers for overwintering and return to *Prunus* in early spring to mate.

**Figure 1.**
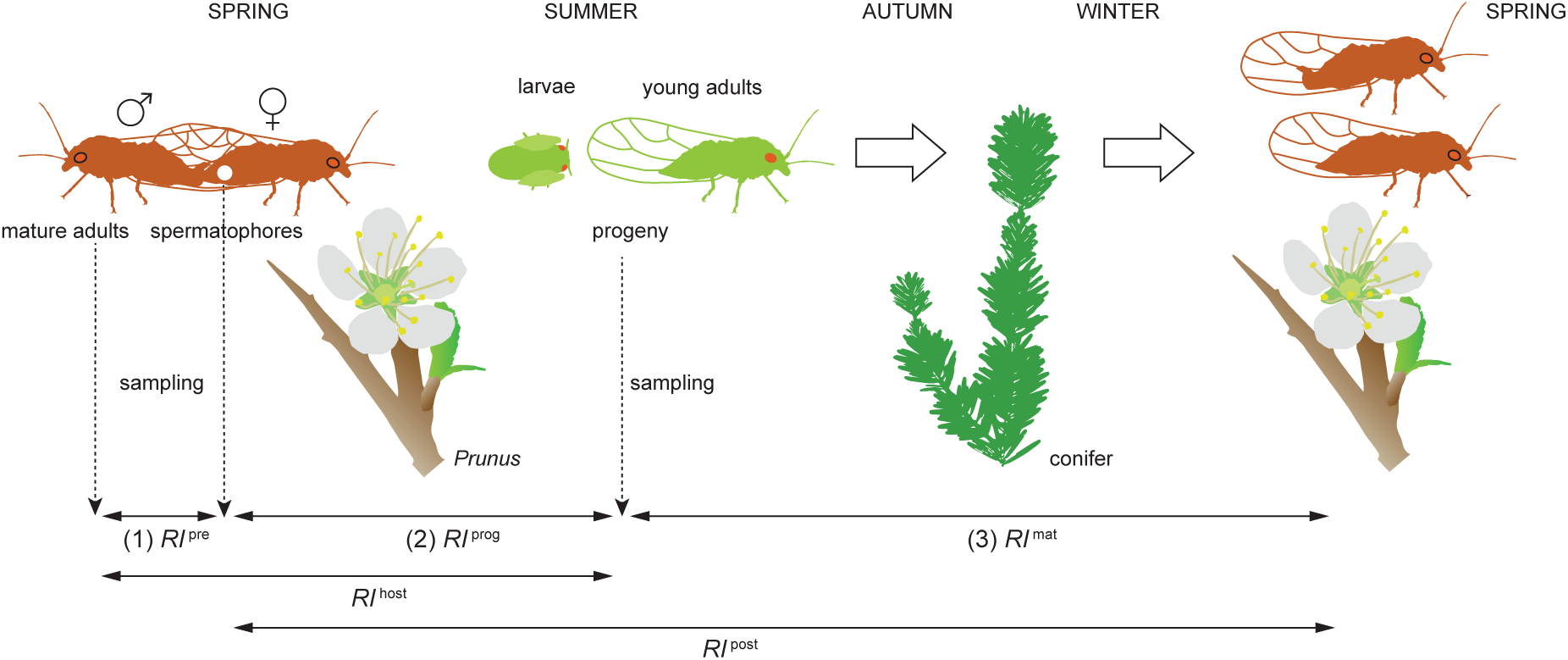
Life cycle of *Cacopsylla pruni* and sampling used to estimate reproductive isolation (RI) between its cryptic species at various barriers, or combinations of barriers. Barriers are shown as horizontal arrows and their effects are estimated with RI indices defined in the main text.

Rearing *C. pruni* in controlled conditions has proven extremely difficult (Jarausch and Jarausch 2016). However, the non-overlapping generations and co-occurrence of the *C. pruni* species at several sites make them good candidates for field-based estimates of RI within their life-cycle (Figure 1). To this aim, we genotyped mating adults, inseminated spermatophores and progeny as species *A* or *B* or as hybrids.

### Sample collection and species assignment

Psyllids were collected in spring 2010 on *Prunus* in southern France at three sites: near Tautavel (42°47′38N, 2°41′56E), Grabels (43°39′35N, 3°49′12E), and Bompas (42°43′43N, 2°56′31E). We also used collections obtained in spring 2008 near Torreilles (42°44′29N, 2°59′6E). Each sampling site consisted of a small number of bushes or hedges of *Prunus* and covered a few dozen meters at most.

Mature adults were sampled at all sites. Progeny (larvae and young adults of the subsequent generation) were sampled at Tautavel and Grabels. Psyllids that fell from beaten branches onto flat nets were either stored in ethanol and/or frozen. Mating pairs caught on nets were stored in separate tubes.

Purification of DNA followed Peccoud et al. (2013). Abdomens of mature females were softened in 70% ethanol for spermatophore extraction. Spermatophores were identified as glossy white pellets in spermatheca under a stereo microscope and transferred separately to DNA purification wells. To minimize between-species DNA contamination risk, each batch of dissections, DNA purifications and amplifications of spermatophore DNA was performed on females of the same species.

Each DNA sample was assigned to species *A* or *B* using a single diagnostic PCR of the Internal Transcribed Spacer 2 (*ITS2*) gene, which yields an amplicon of a specific size for a given species (Peccoud et al. 2013). Individuals showing two bands – putative hybrids – were reprocessed through DNA extraction (re-using their carcasses after washing in water) and PCR in order to minimize the risk of DNA contamination being interpreted as hybridization. To identify the maternal species of each confirmed hybrid, we Sanger-sequenced a mitochondrial region encompassing the *COI* gene (sequences are available under Genbank under accession numbers xxx). Supporting text I details purification, primers, the genotyping of hybrids and possible sources of error.

### Modeling reproductive isolation in Cacopsylla pruni

The genotype data of spermatophores, progeny and mature adults allowed estimation of RI arising: (1) between colonization of *Prunus* by mature adults and insemination, (2) between insemination and the sampling of progeny on *Prunus*, (3) following the sampling of progeny, overwintering on conifers and return of mature adults on *Prunus*. Indices related to (1), (2) and (3) are given superscripts “pre” (for *pre*-*insemination*), “prog” (*progeny*) and “mat” (*mature adults*) respectively (Figure 1).

Gene flow due to inseminations at sampling site 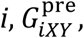 reflects the proportion of spermatophores of species *Y* per female of species *X* (Table 1). All dissected females were inseminated, thus there was no evidence that female reproductive success differed between species. Accordingly, we used equation 8 to model 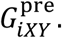 At this stage, gene flow before the barrier reflects species proportions among mating adults of each sex, thus:

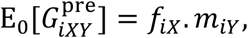

where *f_ix_* and *m_iY_* are the proportions of species *X* and species *Y* among mature females and males of site *i*, respectively. Thus, equation 8 becomes:

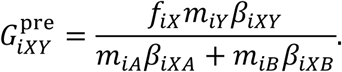

At a subsequent barrier, null gene flow is gene flow through the previous barrier. Thus, from equation 5, gene flows at the two postzygotic barriers are:

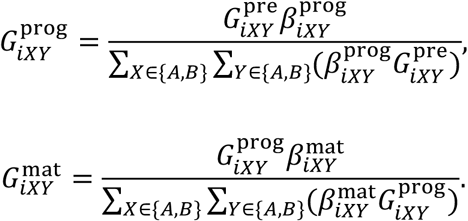

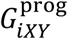 reflects the proportion of *XY* genotypes among progeny, and 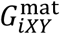 reflects the proportion of *XY* genotypes among returning adults. With these specifications, porosities ***β*^pre^**, ***β*^prog^** and ***β*^mat^** can be estimated from genotype data using Bayesian modelling (below).

Equation 11 was used to compute the combined porosities of successive barriers (Figure 1). Thus, we quantified ***β*^host^** and ***RI*^host^**, representing RI on spring host-plants [barriers (1)+(2)]; ***β*^post^** and ***RI*^post^**, representing global post-insemination RI [barriers (2)+(3)]; and ***β*^total^** and ***RI*^total^** for barriers (1)+(2)+(3). The “absolute contribution” of barriers (Ramsey et al. 2003; Sobel and Chen 2014) was quantified as the difference in cumulative RI either side of each barrier (Ramsey et al. 2003).

### Spatial heterogeneity in pre-insemination isolation

To determine the degree of spatial heterogeneity in pre-insemination RI, ***β*^pre^**, we incorporated a finite mixture model (FMM) (McLachlan and Peel 2000) as a parsimonious model of hidden spatial structure (Pleydell and Chretien 2010). This FMM allocates each study site to one of *k* ∈ {1…*n*} “site groups”, where each site in a group shares identical porosities ***β*^pre^** and *n* is the number of study sites. This introduces vectors ***z***, which allocates sites to groups, ***w***, which weights the importance of groups and ***κ***, an indicator vector that activates / disactivates groups (see supporting text II).

### Bayesian inference

Bayesian analysis of RI in *C. pruni* required making inference from the posterior distribution:

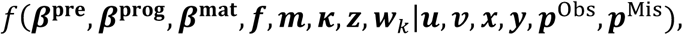

with new terms defined below. Uninformative priors were adopted for all parameters.

Likelihood functions for species frequencies among sexes (***m*** and ***f***) were obtained assuming the number of individuals of each species among sampled males (***u***) and females (***v***) follow multinomial distributions with probabilities ***m*** and ***f*** respectively.

At barrier (1), the likelihood was evaluated using genotype data for inseminated females (***x***) and spermatophores (***y***). The numbers of species *A* and *B* spermatophores extracted from a female of species *X* were assumed to follow a multinomial distribution with probabilities *G_XA_* / (*G_XA_* + *G_XB_*) and *G_XB_* / (*G_XA_* + *G_XB_*), respectively. This assumes independent inseminations – indeed males of the related species *C. pyricola* inseminate one spermatophore per female (Burts and Fisher 1967; Krysan 1990).

At barrier (2), counts of the four genotypes among progeny, ***p*^Obs^**, were assumed to follow a multinomial distribution with probabilities ***G*^prog^**. The likelihood also accounted for two hybrids (from Tautavel) of unknown maternal ancestry, ***p*^Mis^**, that failed to amplify at the mitochondrial region (see supporting text I and II).

The likelihood at barrier (3) was derived similarly to that of barrier (2), from genotype data ***u*** and ***v***, neglecting possible between-year differences in genotype frequency. Further model details, and a glossary defining all variables, are provided in supporting text II.

The posterior distribution was sampled using Markov chain Monte Carlo (MCMC) (Gelman et al. 1995). Site-group activation indicators, ***κ***, were sampled using Reversible Jump MCMC (Green 1995). The model and MCMC algorithm were written and executed in NIMBLE 6.10 (de Valpine et al. 2017) within R 3.4.1 (R Development Core Team 2017). One hundred MCMC chains of 600,000 iterations were run, the first 100,000 iterations were removed as burn-in and samples were saved each 50 iterations. Concatenated output (10^6^ samples in total) was analyzed using R package CODA. Source code and data are available at https://bitbucket.org/DRJP/reproductive_isolation_mcmc/

## Results

Table 2 summarizes the genotypic data and shows large differences in species frequencies across sites.

**Table 2.** Genotype data (assignment to *C. pruni* species *A* or *B*) from mature adults, spermatophores and progeny at the four sampled sites.

|  | Tautavel |  | Grabels |  | Bompas |  | Torreilles |  |
| --- | --- | --- | --- | --- | --- | --- | --- | --- |
|  | *A* | *B* | *A* | *B* | *A* | *B* | *A* | *B* |
| Females | 307 | 87 | 127 | 41 | 30 | 31 | 60 | 146 |
| Males | 211 | 57 | 43 | 18 | 27 | 23 | 26 | 64 |
| Intra-inseminated females | 194 | 76 | 110 | 38 | 30 | 30 | 53 | 141 |
| Cross-inseminated females | 1 | 11 | 1 | 3 | 0 | 1 | 0 | 5 |
| Spermatophores in *A* females | 826 | 1 | 249 | 1 | 80 | 0 | 83 | 0 |
| Spermatophores in *B* females | 14 | 202 | 3 | 65 | 1 | 70 | 5 | 212 |
| Progeny from *A* mothers | 1935 | 3 | 382 | 0 |  |  |  |  |
| Progeny from *B* mothers | 10 | 424 | 2 | 142 |  |  |  |  |

For logistic reasons, not all genotyped females were dissected. “Cross-inseminated” refers to females carrying at least one allospecific spermatophore. For the progeny column headings, *A* or *B*, indicate the paternal species.

We did not model premating isolation as only 46 mating pairs were caught on sampling nets, all at Tautavel. Thirty-five involved individuals of species *A*, and 11 involved individuals of species *B*. No heterospecific pairs were found. Species proportions in mating pairs were indistinguishable from those in mature adults (*χ*^2^ = 0.045, *p* = 0.84, 1 d.f.) but differed significantly from those expected under random mating (*χ*^2^ = 40.7, *p* < 0.001, 1 d.f.). The sampling time of 41 mating pairs sampled over the course of a single day showed little difference between species (Mann-Whitney = 244, *p* > 0.19), providing no evidence for differences in timing of mating activities.

Most spermatophores (1812 of the 1990 extracted) were successfully genotyped (missing data is discussed in supporting text I). Interspecific inseminations were detected at all sites (Table 1) and involved 1.38% of genotyped spermatophores. This indicates strong but incomplete pre-insemination isolation (*RI*^pre^, Figure 2A).

**Figure 2.**
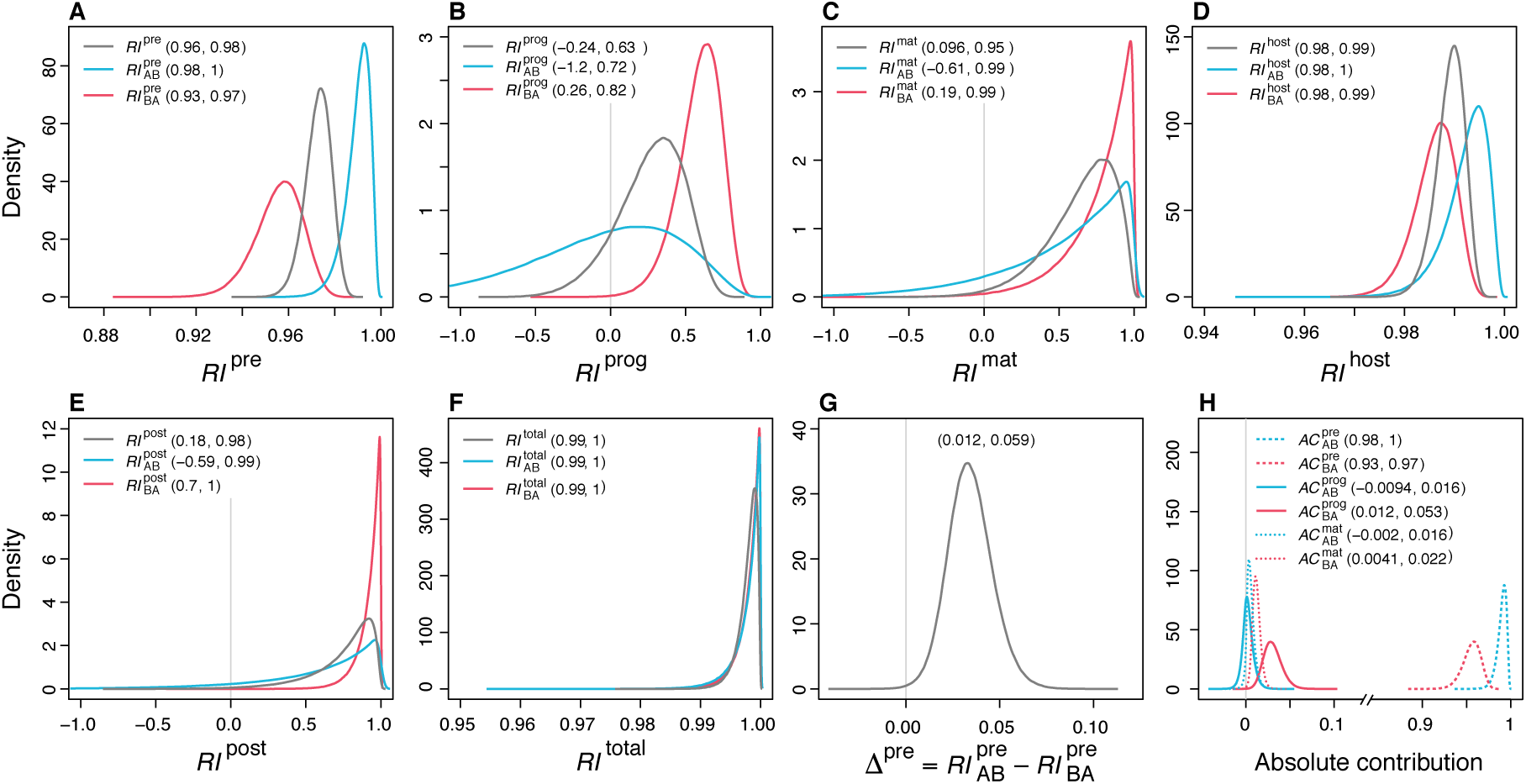
Posterior probability distributions of reproductive isolation (RI) between *Cacopsylla pruni* species measured at three reproductive barriers (panels A, B, C) and their combinations (panels D, E, F); (G) asymmetry in pre-insemination RI; (H) absolution contributions of reproductive barriers to overall RI. Ninety-five percent credibility intervals are shown in parentheses. See Figure 1 for a representation of the different forms of RI measured in *C. pruni*.

The proportion of MCMC samples in which ***RI*^pre^** differed between sites was ~0.001, providing only negligible evidence for between-site variation. In terms of asymmetry, *RI*^pre^ was stronger in *A*×*B* crosses than in the opposite direction, Δ^pre^ being positive (Figure 2G). Other barriers showed little evidence for asymmetry, as posterior distributions of directional RI indices for reciprocal crosses largely overlapped (Δ not shown).

Results support positive post-insemination isolation against *B*×*A* hybrids of the progeny 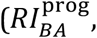 Figure 2B), meaning these hybrids were less frequent than expected from cross-inseminations. The absence of hybrid genotypes in mature adults (Table 2) rendered *RI*^mat^ positive for *B*×*A* hybrids (Figure 2C), indicating mortality between emigration from spring hosts and return to these hosts the subsequent year. For the opposite cross direction, uncertainty was large, due to the strong isolation against *A*×*B* insemination and subsequent low expected frequency of *A*×*B* hybrids.

The combinations of these successive reproductive barriers led to strong *RI*^host^, *RI*^post^ and essentially complete overall RI (Figure 2D-F). Pre-insemination barriers contributed by far the most to overall RI, as shown by the high absolute contribution *AC*^pre^ (Figure 2H).

## Discussion

### Benefits and assumptions of the approach

This work introduces the notion of barrier porosities, which represent contingency-independent probabilities of gene flow, to facilitate RI estimation. Our formulations extend the current RI quantification framework (Sobel and Chen 2014) in several ways.

First, they standardize the construction of RI indices for any type of barrier my modelling null gene flows E_0_[***G***] and potential gene flows ***G*** (Table 1) at each barrier.

In addition, by explicitly considers all four cross-types (within and between species), this approach leads naturally to the construction of directional RI indices (*RI_XY_*, equation 2). These indices share the properties of Sobel and Chen’s (2014) bidirectional *RI* and satisfy the growing interest in measuring asymmetry in RI (Lowry et al. 2008; Matsubayashi and Katakura 2009; Sanchez-Guillen et al.; Yukilevich 2012; Brys et al. 2014). A notable difference with the bidirectional RI indices of Sobel and Chen (2014) and of equation 1 is the asymmetry of *RI_XY_*, which varies from −3 to 1, and not between −1 and 1. However, a value of −3 accurately informs that directional gene flow is 300% higher than expected under random mating (1/4). It therefore seems sensible to prioritize linearity with gene flow (barrier porosity) over symmetry of the RI index.

Our formulation also simplifies the quantification of cumulative effects of sequential and potentially asymmetrical barriers on RI – it is sufficient to compute a normalized product of sequential porosities estimated separately (equation 11). This reduces the difficulty of formulating RI over sequential barriers, where exigent checking for correctness in respect to a particular combination of barriers is typically required (Sobel and Chen 2014). Because barrier porosities are probabilities, and are designed to be contingency-independent, they can be combined (multiplied) for any sequence of barriers studied by any method ranging from field surveys to laboratory experiments (so long as phenotypes controlling RI are not significantly affected by test conditions).

Finally, because our porosity-centered specification permits comparison of modelled and observed gene flow, it readily accommodates Bayesian inference and hence credibility intervals for RI-related indices. The potential of Bayesian modelling is demonstrated here with a finite mixture model designed to detect spatial heterogeneity in RI. These developments can help identify local factors conditioning RI. In particular, were RI to vary according to species frequencies, one may question the two main assumptions underlying frequency-independent RI indices: (i) hybrid survival rates are unaffected by genotype frequencies in the progeny (implied by equation 4), neglecting possible effects of competition on hybrids, and (ii) the risk of hybridization per interspecific encounter is stable (equation 7). While we could not evaluate the former assumption due to uneven sampling of progeny and the scarcity of hybrids, the latter is discussed in the next section.

### Intensity and contribution of reproductive barriers

The Bayesian model used to analyze genotypic data from *C. pruni* populations demonstrated high, asymmetrical pre-insemination isolation (***RI*^pre^**) with little evidence for between-site variation, and positive post-insemination isolation (***RI*^prog^** and ***RI*^mat^**) against *B*×*A* crosses (Figure 2). The combination of these barriers results in essentially complete RI in both directions.

Pre-insemination isolation is dominated by premating isolation, given the absence of heterospecific mating pairs among the 46 collected. Conspecific mate preference could be mediated by olfaction (Soroker et al. 2004; Wenninger et al. 2008; Guedot et al. 2009) and/or acoustic signals (Percy et al. 2006; Tishechkin 2007; Wenninger et al. 2009), both of which contribute to species recognition and mate attraction in other psyllid species. Mechanical isolation (Sota and Kubota 1998; Holwell et al. 2010) appears unlikely, as variation in male genitalia morphology could not be detected by optical and electron microscopy (N. Sauvion, unpublished). The same can be said for temporal isolation [reviewed in Taylor and Friesen (2017)], as the timing of mating did not significantly differ between species according to mating pairs caught within the course of a day. At larger temporal scales, synchrony between reproductive cycles is supported by the similar species proportions across larval stages at Tautavel (*χ*^2^ = 2.0556, *p*>0.35, 2 d.f.).

We detected that pre-insemination RI significantly differs according to the direction of crosses (Figure 2A,G), suggesting that *B* females and/or *A* males are on average less discriminatory than their allospecific counterparts in respect to species recognition. Asymmetric pre-zygotic isolation is frequently observed in mate preference assays (Jaenike et al. 2006; Rafferty and Boughman 2006; Takami et al. 2007; Dopman et al. 2010; Raychoudhury et al. 2010; Merrill et al. 2011; Veen et al. 2011; Sanchez-Guillen et al. 2012), but is only rarely measured in the field (Bournez et al. 2015). In comparison to laboratory studies, the asymmetry we observed involves much higher levels of RI (Figure 2A). This suggests that asymmetry in pre-zygotic isolation can persist late in the speciation process, as does prezygotic RI in general (e.g., Coyne and Orr 1997; Mallet et al. 2007; Merrill et al. 2011), and/or that premating RI can be higher in the field than in the laboratory (Jennings and Etges 2010).

Interestingly, we found no convincing evidence that pre-insemination RI varied between the four sampling sites, despite large differences in relative species frequencies (Table 1). Hence, the assumption of a stable hybridization risk per interspecific encounter (implied by equation 7) is not called into question. Keeping in mind that our ability to challenge this assumption is limited by the number of sampling sites, our observations inform us on the bases of incomplete prezygotic RI in *C. pruni*. A stable risk of mating per interspecific encounter may indicate a certain degree of conspecific mate preference that is both relatively insensitive to site-specific factors, and similar among individuals of the same species and sex (e.g., Merrill et al. 2011). The hypothesis of between-individual variation in mate preference, potentially due to polymorphism at underlying loci, would less parsimoniously explain incomplete RI. Indeed, it would not explain why cross-inseminations appear more frequent at species *A*-rich sites, unless the less discriminatory individuals essentially occurred among males of species *A*. This hypothesis thus also requires that mate choice be mostly exercised by males. Mate-choice experiments would help to evaluate these hypotheses.

The predominant contribution of prezygotic barriers to overall RI (Figure 2H) naturally follows from their early occurrence in the species life-cycle and has been reported in various sympatric species (Ramsey et al. 2003; Malausa et al. 2005; Kay and Husband 2006; Lowry et al. 2008; Matsubayashi and Katakura 2009; Sanchez-Guillen et al. 2012). This predominance does not indicate that post-insemination barriers contributed little to the divergence of *A* and *B* species. These barriers may have reinforced premating isolation [see Servedio and Noor (2003); Coyne and Orr (2004) for a review on reinforcement] and have certainly permitted genetic divergence between *C. pruni* species (Sauvion et al. 2007; Peccoud et al. 2013) in the face of cross-insemination. Some barriers (***RI*^prog^**, Figure 1) operate between insemination and progeny growth, at least for *B*×*A* crosses (Figure 2B), and others (***RI*^mat^**, Figure 2C) affect survival of grown hybrids up to their return on *Prunus* shrubs. In terms of causes, ***RI*^prog^** possibly reflects low sperm efficacy in allospecific females (e.g., Matute 2010) and/or reduced hybrid survival up to sampling.

Although the scarcity of hybridization in *C. pruni* limits the precision of certain estimates, our case study illustrates how the proposed framework provides estimates of reproductive barriers at an arbitrary number of sampling points through the species life cycle. Future models could incorporate refinements such as independent development and/or survival rates for each sex and developmental stage, more sophisticated models of spatio-temporal variation, or other sources of prior information.

## Acknowledgements

We thank René Rieu for his advice on spermatophore extraction, and to Gaël Thébaud, Gérard Labonne and François Bonnot for helpful comments on the study and drafts of this article. We thank Josiane Peyre and Patrick Limon for their help in genotyping. This work utilized computing resources form INRA’s MIGALE cluster (http://migale.jouy.inra.fr) and benefitted from the assistance of Eric Montaudon and Véronique Martin. Part of this work was funded by INRA grant SDIPS (Speciation and molecular Diagnosis of Insect Pest Species complexes).

## Conflict of interest

The authors declare no conflict of interest.

